# Identification of *Pectobacterium* species isolated from the soft rot of tetecho (*Neobuxbaumia tetetzo*), a columnar cactus, and associated metagenomics

**DOI:** 10.1101/2021.02.01.429127

**Authors:** David Vargas-Peralta, Delia A. Narváez-Barragán, Andrés de Sandozequi, Miguel F. Romero-Gutiérrez, Lorenzo Segovia, Claudia Martínez-Anaya, Luis David Alcaraz, Rodolfo de la Torre Almaraz

## Abstract

*Neobuxbaumia tetetzo*, commonly known as tetecho, is a columnar cactus endemic to Mexico. In the last 15 years, damage has been observed in young and adult plants of *N. tetetzo*, ranging from chlorotic spots with a wet appearance in early stages to tissue necrosis in advanced stages and finally the death of the plant; *Pectobacterium brasiliense* is the causal agent of the damages. Disease progression may be delayed or accelerated by the involvement of other bacteria, either pathogenic or endophytic, at the site of infection. Our goal was to confirm the presence of *Pectobacterium brasiliense*, in the soft rot of *N. tetetzo* and to determine the presence of other bacteria associated with the rot. We isolated three bacterial strains (A1, A3 and A8) from diseased tissue from three separate *N. tetetzo* plants, and compare them using biochemical and molecular techniques, such as whole-genome sequencing of strains A1 and A3. Phylogenetic analyzes confirmed that A1 corresponded to *P. brasiliense*, whereas A3 was more misimlar to *P. polaris*. Additionally, sequencing of 16S rRNA gene from metagenomic DNA isolated from healthy and diseased tissue of *N. tetetzo* indicated the presence of four operational taxonomic units (OTUs) at the order level, unique to the diseased tissue: Actinomycetales, Burkholderiales, Caulobacterales, and Sphingomonadales, with probable participation in the soft rot process.

## Introduction

The Tehuacán-Cuicatlán Valley, Mexico, is a semi-arid zone covering an area of almost 10,000 km^2^, it is considered one of the dry areas with the greatest diversity of flora and fauna, in addition to hosting a large number of endemic species. One endemic group of plants that stand out are the columnar cacti (Pachycereeae tribe) [1,2]. Zapotitlán de las Salinas is a community within the valley (18°20’N, 97°28’W), which houses the endemic columnar cactus *Neobuxbaumia tetetzo*, commonly called tetecho or teteche, forming dense forests of up to 1,200 individuals per hectare known as tetecheras. These tetecheras are of outmost importance for local inhabitants, since they are used as living fences, construction materials, firewood, food and ornamental plants, and are also of ecological relevance since they interact with different types of pollinators [3-5]. Observations of the last 15 years document plant damage and disease as follows: a) development of wet-looking chlorotic spots that may appear at the base, in the middle or in the upper part of the plant; b) spots turn gray; c) in advanced stages of the disease the tissue cracks and exudes a brown fluid; d) flesh detachment occurs exposing the woody tissue, and what is left dries out acquiring a dark gray aspect; eventually the plant may collapse (Fig 1).

**Fig 1.**
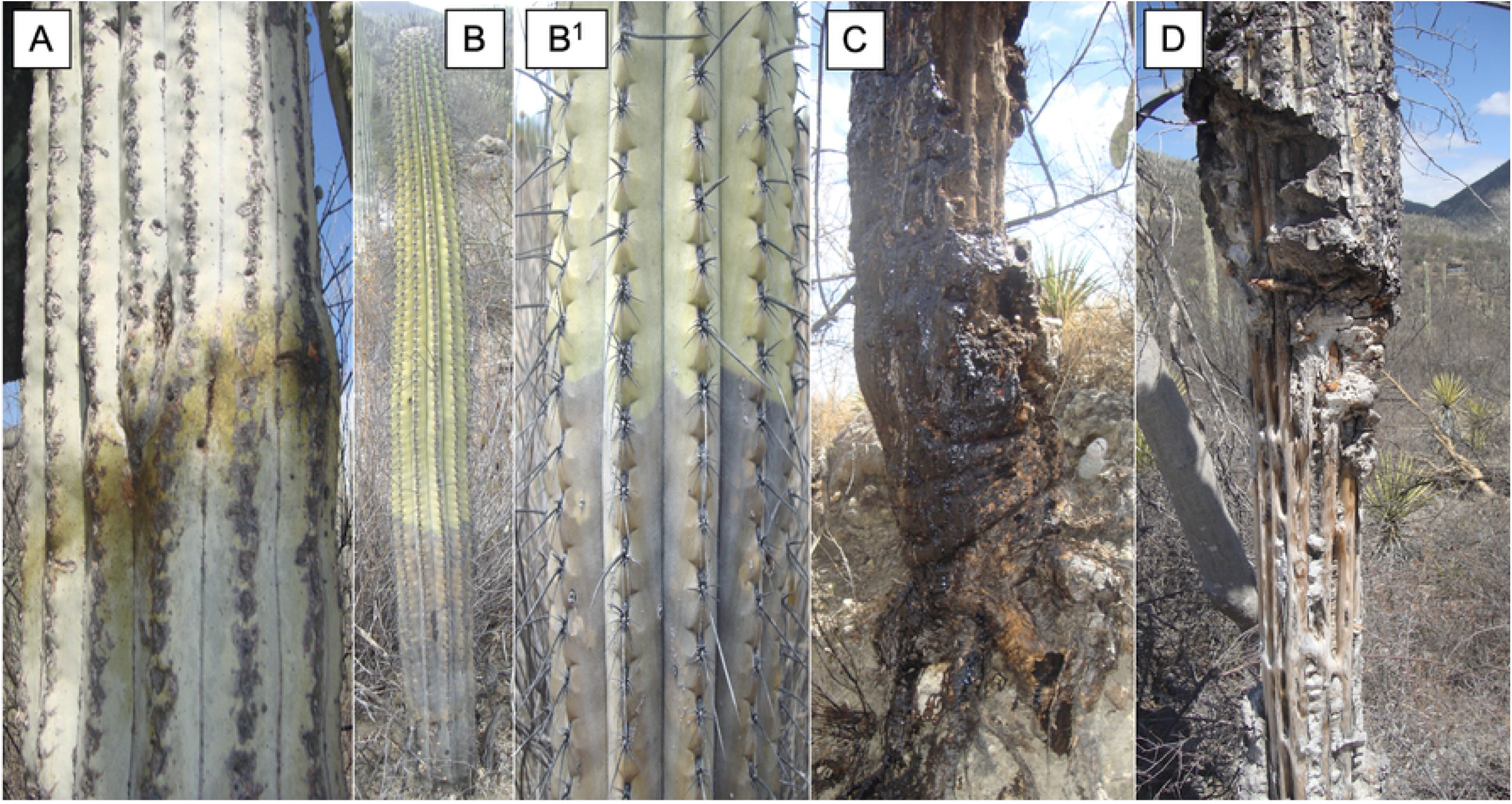
Progression of soft rot in *N. tetetzo.* The onset of rot begins with wet-looking chlorotic spots (A); these spots turn gray (B and B^**1**^); in advanced stages tissue loss and brown exudates occur (C); finally, the plant dries out and dies (D).

Damage is mainly attributed to the phytopathogenic bacteria *Pectobacterium carotovorum* subsp. *brasiliense* (now to elevated to the species *P. brasiliense*), identified by MejÍa-Sánchez and collaborators (2019). The authors characterised 10 out of 80 isolates using pathogenicity and biochemical tests, comparison of the partial sequence of the 16S rRNA gene against the nt database of GenBank, amplifing the intergenic spacer region (ISR) 16S-23S rRNA with the BR1f / L1r oligonucleotides [7], and phylogenetic reconstruction using multiple alignments of the sigma factor 38 of the RNA polymerase subunit (*rpoS*). Although analysis of the 16S rRNA gene and the *rpoS* gene are a first aproach for species identification [8], more information can be used for better taxonomic resolution either at the species or subspecies level. Although the region amplified by the BR1f / L1r oligonucleotides is specific for *P. brasiliense*, it lacks evidence to confirm the presence of the *P. brasiliense* in *N. tetetzo*, but provides a first approximation.

Disease progression may be delayed or accelerated by the involvement of other bacteria, either pathogens, or endophytes that under certain conditions are able to participate in the development of symptoms [9][10]. The use of classical microbiology techniques coupled with Next Generation Sequencing (NGS) technologies to obtain partial or complete sequences of bacterial genomes, allows better characterization of new bacterial isolates from *N. tetezo* providing better support and resolution for bacterial taxonomy [8]. NGS technologies also aid in the analysis of microbial populations and communities [11], to understand the participation of two or more bacteria and their contribution to the occurrence of plant diseases. We aimed to confirm the presencie of *Pectobacterium brasiliense* during *N. tetetzo* soft rot and to explore the bacterial diversity associated with the pathology of this impostant cactus.

## Materials and methods

### Tissue collection and isolation of *Pectobacterium* sp

Two samples of healthy and diseased tissue from *N. tetetzo* from the Zapotitlán de las Salinas region were collected. Potatoes were used as bait in the first sampling to isolate the primary causative agent, *Pectobacterium* sp. A total of four potatoes were inoculated, three with diseased tissue and one with healthy tissue (control). Rotten potato tissue was scraped and streaked in Petri dishes with nutrient agar medium (Merk), incubated at 37 ° C for 48 h, several passes were performed under the same conditions until isolated colonies were obtained. Three *Pectobacterium* sp isolates were obtained, and assigned the names A1, A3 and A8. In the second sampling, healthy and diseased tissue from *N. tetetzo* was taken with a perforation tool and placed in sterile centrifuge tubes and transported to the laboratory.

### Phenotypic characterization

Biochemical tests were performed according to Schaad et al. (2001) for the identification of phytopathogenic bacteria, including: Gram stain, KOH test, anaerobic growth in Hugh and Leifson medium, mucoid colonies in yeast extract-dextrose-CaCO3 (YDC) medium, yellow or orange colonies in YDC and Nutrient-broth yeast extract agar (NBY) and fluorescence in King B medium (KB). Three repetitions per isolate (A1, A3 and A8) were performed for each test.

### Pathogenicity tests and host range

Plant material was previously disinfected with 5% sodium hypochlorite and rinsed with distilled water. From fresh Petri dish cultures of the three isolates, colonies were taken and diluted in phosphate buffer pH 7.4 at OD_600_=0.4 (1×10^32^ cells), each *N. tetetzo* plant of approximately four years old was inoculated with 3-ml syringes. Host range was tested in other Cactaceae species: *Opuntia ficus-indica* cladodes (prickly pear) and *Pachycereus pringlei* (elephant cactus). Six squares (2 cm each side) were traced on cladodes of *Opuntia ficus-indica*, four of them were inoculated with 500 µL of the bacterial suspension and the remaining two were inoculated with 500 µL phosphate buffer and sterile H_2_O as controls. Cladodes were incubated under greenhouse conditions for 17 days. *P. pringlei* were inoculated by duplicate, in a small cut with 10 µl bacterial suspension (1×10^8^ cells) per isolate from overnight cultures in LB medium and incubated in a humid chamber at 30°C for 24 h.

### Determination of virulence

Virulence was determined as the extent of plant tissue degradation and quantified by tissue loss on celery stalks. Five-mm cuts on disinfected (with 5% sodium hypochlorite for 15 min) celery stalks pieces of approximately 15 cm, were inoculated with 10 µl of diluted to OD_600_=0.1 the different isolates overnight cultures in LB medium (n=33 per strain). These were incubated in a humid chamber at 30°C for 24 h. Tissue loss was determined by the difference of weight before and after removing the macerated tissue.

### Detection of plant cell wall degrading enzymes (PCWDE’s)

Enzymatic activity was detected in plate assays. LB agar medium (Bacto) was prepared with 2% sodium carboxymethyl cellulose (CMC) (Merck) or 2% pectin (Agdia AG366) prepared according to the provider instructions; bacteria were plated 30 min after pouring the media. A drop of 3 µL of each strain diluted to OD_600_= 0.1 was placed in the center the Petri dish on both substrates (by triplicate), allowed to dry for 20 min and incubated in a humid chamber at 30°C for 48 h. Enzymatic activity towards CMC was revealed with Congo red 1% for 15 min, then rinsed with 1 M sodium chloride. Pectinolytic activity was detected with 0.1% toluidine blue for 10 min and washed with 20 mM HEPES pH 7.5. Enzymatic index (EI) was calculated by dividing the hydrolysis halo by the colony halo [13].

## Identification of strains A1, A3 and A8

### End-point PCR targeting of the 16S rRNA gene

Genomic DNA was extracted using the modified method of Dellaporta et al. (1983), for PCR (Taq DNA polimerase recombinant, Invitrogen) reactions with oligonucleotides FD1 (5’ AGAGTTTGATCCTGGCTCAG 3’) and RD1 (5’ AAGGAGGTGATCCAGCC 3’) for a 1,600 bp amplicon [15]. PCR set up was: 94°C / 3 min, 30x (94 ° C / 1 min, 55°C / 30 s, 72°C / 2 min) and 72°C / 7 min. PCR products were purified (Wizar SV kit, Promega), and sequenced (at the FESI-UNAM Molecular Biochemistry Laboratory), for comparison against the non-redundant nucleotide database (Nucleotide collection, nr/nt) of the GenBank (NCBI), using BLASTn [16,17], with Megablast program, with a cut-off value of E from 1 to 6, a word size of 28 and a mismatch score of -2.

### Genome sequencing, annotation and comparative genomics

Strains A1 and A3 obtained from the first field sampling were selected for genome sequencing at the UUSMD, Instituto de BiotecnologÍa-UNAM. Sequencing quality was analyzed with the free software FastQC High Throughput Sequence QC Report, version 0.11.7 https://www.bioinformatics.babraham.ac.uk/projects/download.html. Genome were assembled with Velvet version 1.2.10 [18], using a sweep (33 > K < 53) to select the best K value for each draft genome. K-value was selected with QUAST program [19]. Draft genomes were annotated with Prokka v1.11 program [20]. Venn diagram was constructed as previously described in [21]. Predicted proteins of strains A1 and A3 were compared to reference genomes of *P. brasiliense* BC1 (NZ_CP009769.1) and *P. polaris* NIBIO1392 (NZ_CP017482.1), and against each other (all-vs-all BLAST) with Proteinortho v5.16 using reported parameters (p-value <10^−5^, identity >35%, and match lengths >75%) [22]. Subsystem assignment, circular genome construction, and virulence, transport and antibiotic resistance genes were predicted with PATRIC suit [23].

### Phylogenetic analysis

For comparison genomics two analyzes were used: Pyani python average nucleotide identity (ANI) module using the BLAST algorithm (ANIb) [24], with 100 *Pectobacterium* genomes (available until march of 2018, S1 Table), with a suggested cut-off value of 95-96% to delimit bacterial species; the second analysis was based on coding sequences (CDS) of 120 genomes (113 genomes of *Pectobacterium* sp and 5 genomes of the genus *Yersinia* (available until october of 2018; S1 Table) for the construction of a phylogenetic tree, for this the distance measurement between species was established with a Genomic Similarity Score (GSS) based on the sum of the bit scores of shared orthologs, detected as Reciprocal Best Hits (RBH), normalized against the sum of the bit scores of the genes compared against themselves (scores of own bits). The GSS varies in values from 0 to 1, where all the orthologous proteins between two proteomes that are identical has a maximum value of 1, on the contrary, when two proteomes do not show similarity in orthologous proteins, it has a value of 0 [25], the GSS matrix was converter using the expresing 1-GSS, from this, a distance matrix of the GSS scores of the shared orthologs for the proteomes used in a Neighbor-Joining tree is drawn to visualize the genomic distance between the isolates; the tree was built from 1,000 replicas [26].

### Massive sequencing of amplicons of the 16S rRNA gene from samples of *N. tetezo*

For a second sample collected, metagenomic DNA was extracted using the DNAzol^®^ Reagent kit (Invitrogen) from healthy and diseased tissue from *N. tetetzo*, and massive sequencing of the 16S rRNA gene of the V3 and V4 hypervariable regions was performed at the UUSMD Unit-IBT, UNAM, by pair-end sequencing in an Illumina NextSeq® 500 sequencer. For the processing of the sequences, the pipeline was followed according to the Standardized Operating Protocols (SOP) Environmental Genomics FC-UNAM (http://genomica.fciencias.unam.mx/github/). Briefly, the quality of the sequences was visualized with the FastQC High Throughput Sequence QC Report software, version 0.11.7. [27]; quality control and alignment of the readings were performed using the CASPER software [28]. With QIIME program, *subsample_fasta*.*py* random sequences were taken from the files to be analyzed. Grouping and selection of Operational Taxonomic Units (OTUs) were performed using the CD-HIT-EST [29] program with an identity cut-off 97%, and representative sequences were selected with the option *pick_rep_set*.*py* from QIIME. Sequence filters used a first matching against the Greengenes database (gg_13_8_otus; 70% OTUs) eliminating sequences without 16S matches. Matched sequences (70% OTUs) were the input of taxa annotation (97% OTUs Greengenes). The OTUs table was constructed using the QIIME *make_otu_table*.*py* option, chimeras were identified and eliminated with ChimeraSlayer [30-32] mitochondrial, and chloroplast sequences were also identified and removed. Finally, graphs were constructed using R version 3.5.3 sofware with the help of the Phyloseq data package [33].

## Results

### Isolation and Phenotypic characterization of *Pectobacterium sp*. from *N. tetezo*

We have preoviusly reported that *P. brasiliense* causes soft rot in *N. tetetzo* [6], but here we aimed to analyse new isolates in more detail. Using potatoes as baits to collect samples of three diseased and healty individuals we confirmed rot of potatos inoculated with infected cactus tissue while potatoes inoculated with healthy tissue did no rot. Three strains were isolated and designated as A1, A3 and A8 presented the following phenotype: they were negative for Gram staining and positive for the KOH test, thus they are Gram-negative bacteria. All are facultative anaerobic bacteria due to turning Hugh and Leifson medium from blue to yellow. None formed mucoid colonies at 30°C in YDC medium, nor formed yellow or orange colonies in YDC and NBY, respectively, and did not fluoresce in KB medium. Contrarily, all isolates grew well in pectin-containing medium.

### Pathogenicity tests and host range

To determine if the isolates caused sooft rot, young *N. tetetzo* plants were inoculated with each strain. Indeed, all the strains reproduced the symptoms observed in wild cacti: on day one, infected plants presented a slight loss of turgidity, the tissue cracked and exuded a brown fluid. On day two, loss of turgor was more noticeable, the tissue began to darken and the brown liquid continued to emerge, these symptoms worsened as time went on. On day fifteen, inoculated plants lost turgidity, developed a gray-green color and desiccated, indicating their death (Fig 2). These results confirmed that our strains do correspond to the etiological agent of sooft rot.

**Fig 2.**
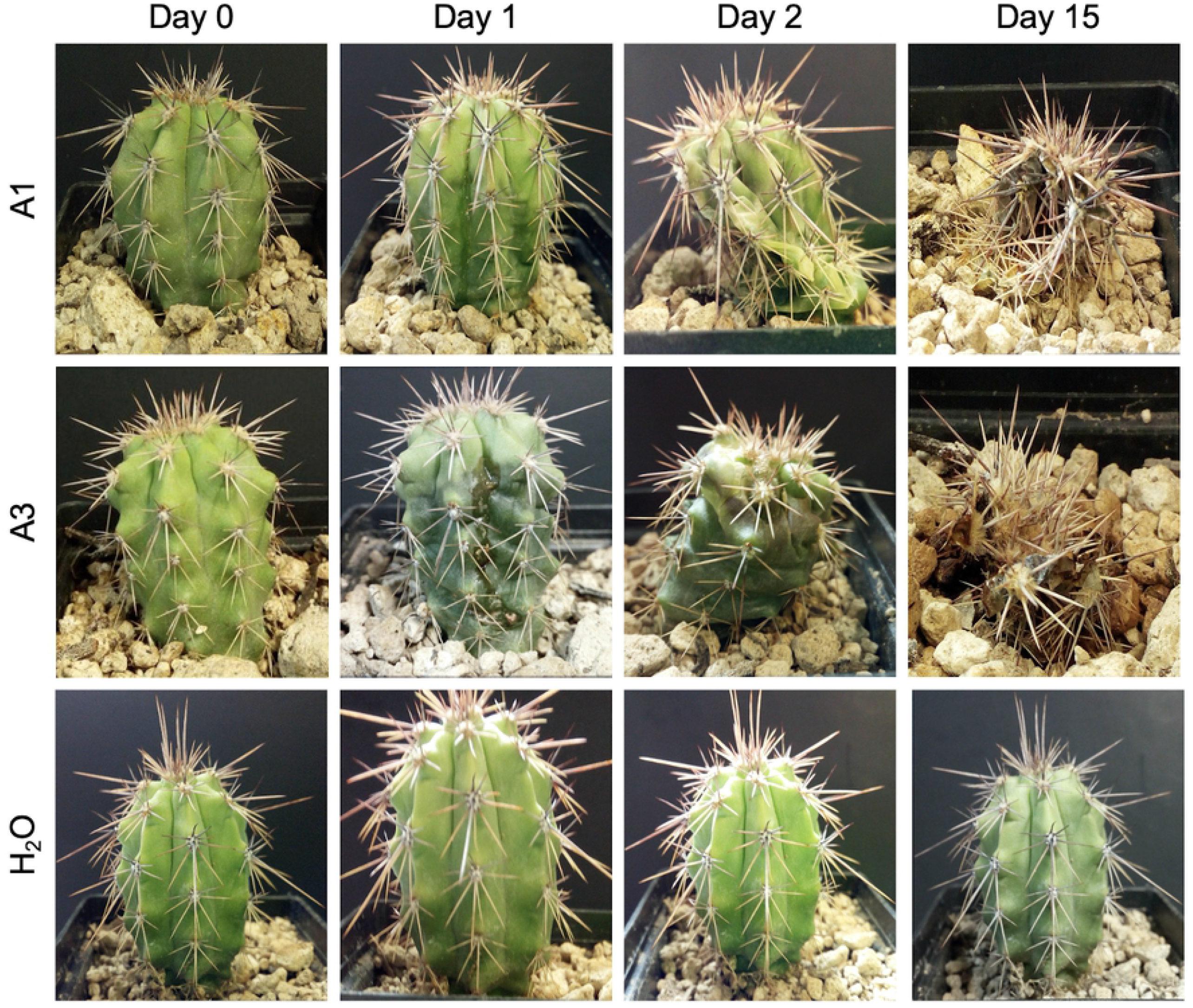
Progressive damage of *N. tetetzo* inoculated with strains A1, A3 and A8. The time from inoculation to plant death is indicated at the top. On the left side the name of the isolates and the control. Only strains A1 and A3 are presented. Top row shows cactus inoculated with strain A1. The second row corresponds to cactus inoculated with strain A3; both isolates caused loss of turgidity, liquid runoff, tissue darkening, desiccation and plant death in a similar period. Third row is a control cactus inoculated with sterile H_2_O that did not show damage during the experiment duration.

We then analyzed whether our strains provocked disease only in *N. tetezo* or whether they are virulent in other cactaceous species: *Opuntia ficus-indica* and *Pachycereus pringlei*. Inoculation of strains in *O. ficus-indica* cladodes developed a wet-looking chlorotic spot at 6 days post-inoculation (dpi); at nine dpi chlorotic spots turned brownish-green, with a brwon necrotic margin; by 12 dpi the spots turned brown, and finally at 17 dpi they darkened, some of spots cracked and exuded a dark fluid. The section inoculated with sterile H_2_O was intact for the duration of the expetiment. Again, the symptoms provoked in *O. ficus-indica* were similar to those observed in the field in *N. tetetzo* (Fig. 3). Finally we analyzed infection of *P. pringlei*. Because individuals of this species were quite small, the bacteria inoculum size was smaller than that used in the previous experiments; this allowed observing differences of virulence between isolates: A1 and A8 showed tissue maceration at one dpi, and at 2 dpi infected plants already showed signs of severe rot and total loss of turgor; meanwhile plants inoculaed with isolate A3 showed a delay of tissue maceration in comparison to A1 and A8, as rot was detected only at 2 dpi. Control plants inoculated with LB medium remained healthy during the time of the experiment (Fig 4). Taken together, these results confirm that strains A1, A3 and A8 recapitulate *in vitro* the disease symptoms observed in wild tetecho cacti, and they point to *Pectobacterium* as the ethiological agent, in agreement to previous observations [6].

**Fig 3.**
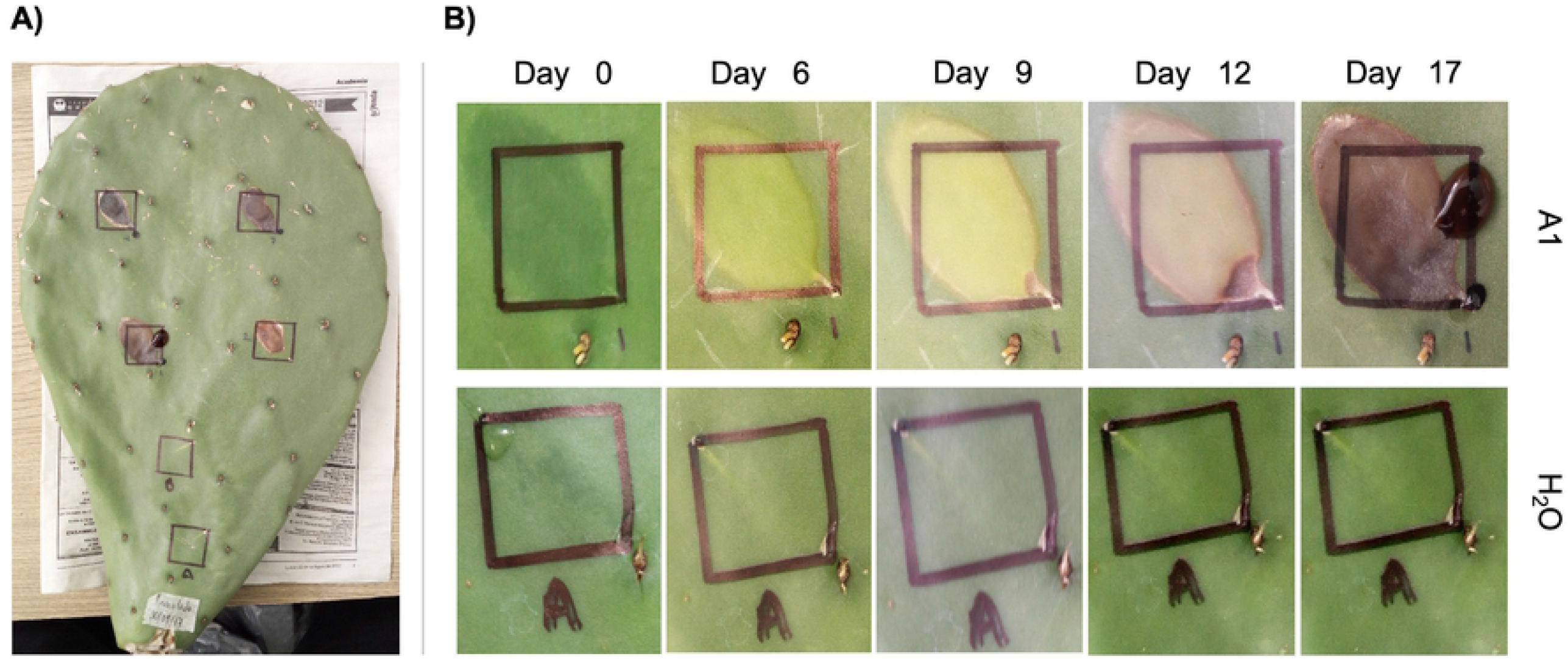
Progressive damage to *Opuntia ficus-indica*. Only isolate A1 is presented. In panel (**A**), prickly pear cactus cladode shows all 6 (2×2 cm) squares, four of which were inoculated with strain A1 and the two remaining squares were inoculated with phosphate buffer pH 7.4 and sterile H_2_O, respectively. In panel (**B**), the top row shows the appearance of symptoms of one infected area through the time, in comparison to the bottom control treatment (with H_2_O) indicated on the right.

**Fig 4.**
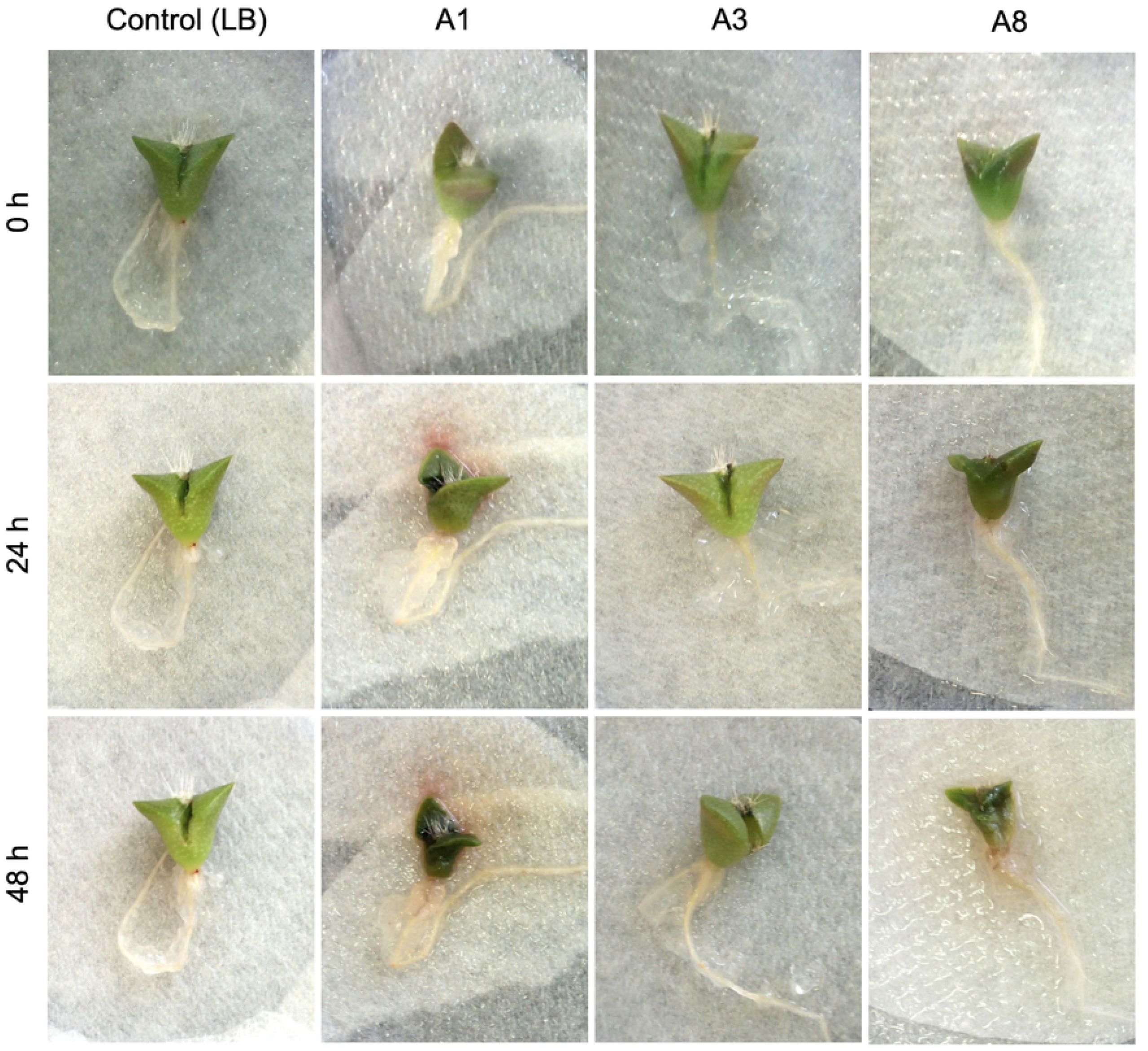
Progressive infection symptoms in *P. pringlei*. Control treatment left is compared with strains A1, A3 and A8 inoculation, through the time (top to bottom).

### Virulence determination and PCWDEs

Given the previous result showing differences between strains, we aimed to compare their virulence in more detail. We used celery stalks as a laboratory model, given its availability and fast symptoms development after inoculation with *P. brasiliense* in comparsion to cacti [34]. We quantified virulence as the amount of tissue loss due to degradation. We found that, similarly to the observed phenotype in *P. pringlei*, isolates A1 and A8 produced more extensive rot in celery (20%) with respect to the isolate A3 (5%) under the same experimental conditions (Fig 5), confirming a different capacity to degrade the plant material between strains. Plant degradation by *Pectobacterium* is due to different plant cell wall degrading enzymes (PCWDEs), thus we asked whether the enzynatic activities between strains also differ. Using a plate assay, we found that the three strains produced halos of comparable size in CMC- and pectin-containing media (Table 1) suggesting that the difference in virulence between strais is independent of their degrading capacity, although it is necessary to analyse individual enzymes produced during infection and other virulence factors.

**Table 1.** Plaque enzyme activity.

| Substrate | Strain | Enzyme index (EI) in cm |
| --- | --- | --- |
| Carboxymethyl cellulose (CMC) | A1 | 2.67 |
|  | A3 | 2.5 |
|  | A8 | 2.67 |
| Pectin | A1 | 2.47 |
|  | A3 | 2.33 |
|  | A8 | 2.47 |

**Fig 5.**
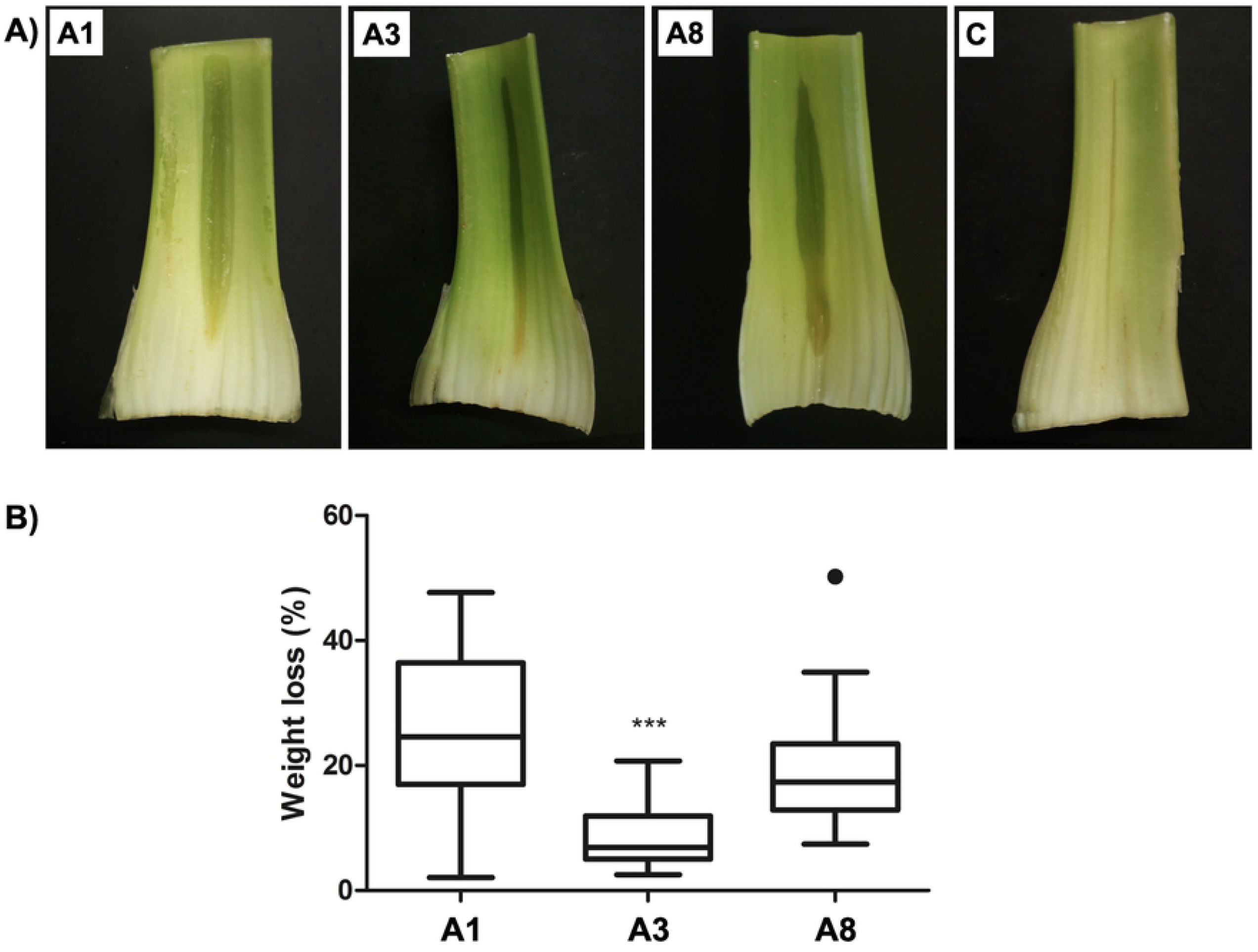
Infection phenotypes in celery. Panel (**A**), phenotypes of infection by strains A1, A3 and A8 compared to inoculation with LB medium control “C”. Panel (**B**), quantification of tissue loss in celery caused by each strain. \**p* ≤ 0.005.

## Molecular characterization of strains A1, A3 and A8, phylogenetic analysis and comparative genomics

### Molecular identification

To confirm the identity of strains A1, A3 and A8 we first analysed the 16S rRNA gene sequence, indicating the following identities: A1 (MG008909.1) and A8 (MG008910.1) were 97% identical to *P. brasiliense* and 100% *P. carotovorum*, respectively. While A3 (MN216189.1) was 98% identical to *P. polaris* and 100% identical to *P. brasiliense*. For a more precise identification we obtained the draft genome of strains A1 and A3 (S1 Figure) (data avaliable in the bioproject PRJNA669375). The two genomes were assembled and annotated (Table 2).

**Table 2.** Data generated from the assembly and annotation of the isolates.

| Feature | Isolates |  |
| --- | --- | --- |
|  | A1 | A3 |
| Total number of contigs | 381 | 326 |
| Largest contig | 124 352 | 263 985 |
| N50 | 46 958 | 59 089 |
| L50 | 32 | 26 |
| % GC | 52.31 | 51.96 |
| Genome size in bp | 4 500 321 | 4 873 548 |
| Coding sequences (CDS) | 4 013 | 4 334 |
| tRNA | 32 | 36 |
| rRNA | 3 | 3 |
| Transfer-messenger RNA (tmRNA) | 1 | 1 |
| Hypothetical proteins<sup>P</sup> | 736 | 859 |
| Proteins with functional assignments<sup>P</sup> | 3715 | 3922 |
| Proteins with EC number assignments<sup>P</sup> | 1107 | 1123 |
| Proteins with GO assignments<sup>P</sup> | 918 | 932 |
| Proteins with Pathway assignments<sup>P</sup> | 812 | 815 |
| Antibiotic Resistance<sup>P</sup> | 44 | 42 |
| Virulence Factor<sup>P</sup> | 46 | 59 |

<sup>P</sup>Predicted with PATRIC suite

### Phylogenetic analysis

According to the ANI analysis, the similarity value between the isolates obtained from *N. tetetzo*, A1 vs A3 was 93.9% below the cut-off value 95-96%, so they are different species (S2 Table). Isolate A1 had an average of 97.1% similarity when compared against 29 *P. brasiliense* genomes, above the cut-off value. When compared to 5 *P. polaris* genomes and 9 *P. carotovorum* genomes, A1 had an average similarity value of 93.9% and 93.3% respectively (S2 Table). Isolate A3 had an average of 96.7% similarity when compared against 5 *P. polaris* genomes, above the cut-off value. When compared against 29 genomes of *P. brasiliense* and 9 genomes of *P. carotovorum*, it had an average similarity value of 94% and 93.3% (S2 Table), respectively (Fig 6). By using another metric such as the genomic similarity score tree (GSS), the tree in this analysis showed three well-defined groups (Fig 7). Group 1 is formed by *Pectobacterium fontis*, a distant species from other *Pectobacterium* because it is shared as a sister clade of groups 2 and 3. Group 2, including *P. betavasculorum, P. wasabiae, P. parmentieri, P. peruviense, P. atrosepticum*, and *P. punjabense*, formed supported and unique subgroups. Finally, Group 3 formed a supported group, with a subgroup including *P. versatil* sp. nov., and *P. odoriferum* sp. nov.; *P. polaris* and *P. brasiliense* are sister groups. To our surprised strain A1 grouped with *P. brasiliense* while strain A3 clustered with *P. polaris*. These results (ANI and GSS) confirmed the identity of *P. brasiliense* for strain A1, and revealed the identity of strain A3 as *P. polaris*, possibly explaining the differences observed by biochemical and pathogenicity tests of each strain.

### Comparative genomics

We then compared A1 and A3 with the reference strains *P. brasiliense* BC1 and *P. polaris* NIBIO1392, with which they are grouped. We found that the four strains share a nucleus of 3,724 coding sequences (CDS). Strains A1 and A3 shared 108 CDSs, most of which are associated with membrane transport. Strain A1 contains 343 unique CDS that participate in membrane transport, energy and generation of metabiolite precursors. Compared with the reference strains, A1 shared 192 CDS with *P. brasiliense* BC1, and 14 CDS with *P. polaris* NIBIO1392. Strain A3 contains 288 unique CDS, mostly associated with membrane transport. And the comparison to the reference strains showed that A3 shares 55 CDS with *P. brasiliense* and 360 CDS with *P. polaris* (Figure 8). Again, this results indicate that A1 and A3 are different species.

**Fig 6.**
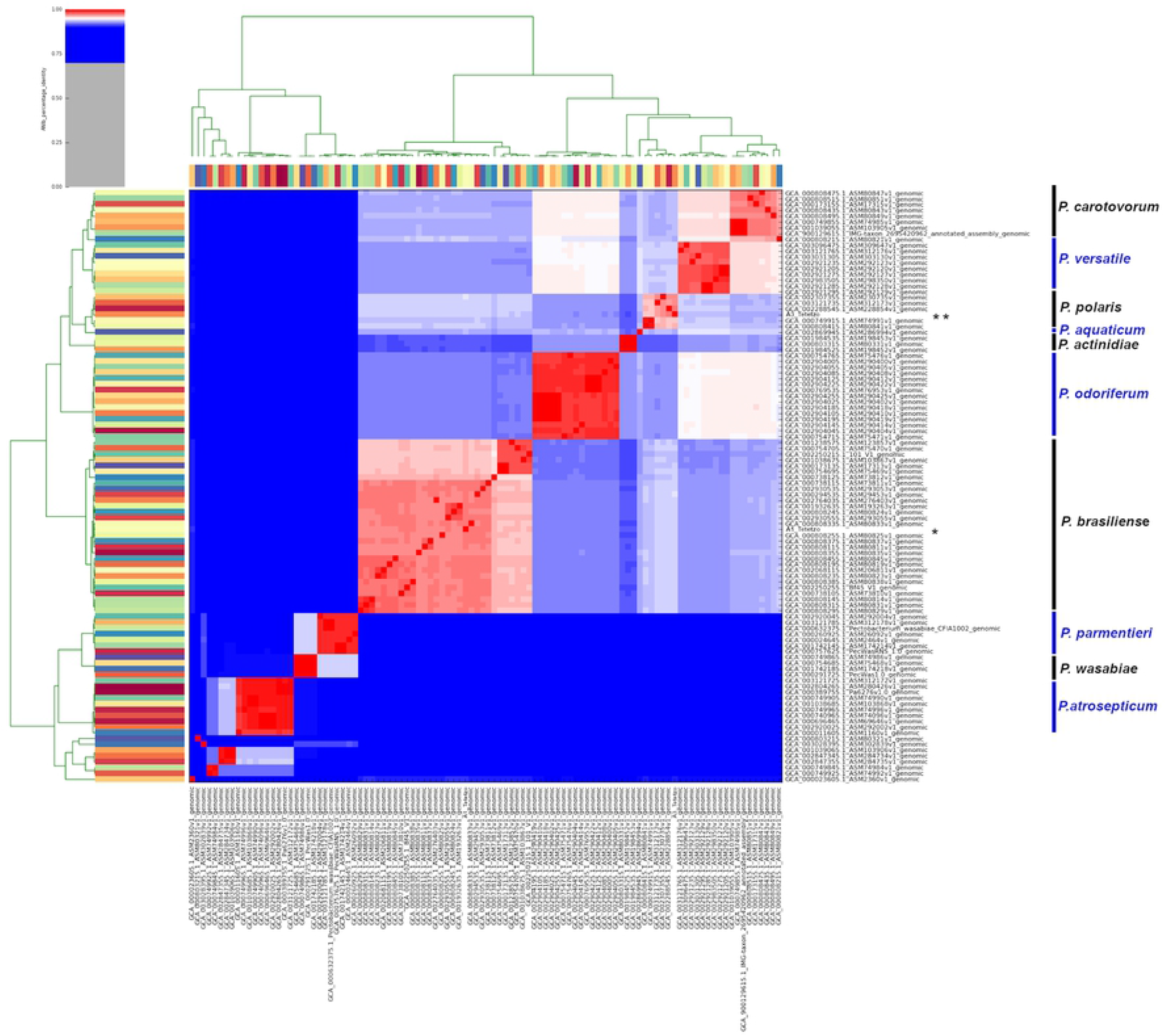
Heatmap of ANI percentage identity for isolates of *Pectobacterium* strains. Heatmap corresponding to 95% ANIb sequence identity are colored red. Blue cells correspond to ANIb comparisons of organisms that do not belong to the same species. Species in the groups formed is indicated on the right side. One asterisk indicates the position of strain A1, and the double asterisk shows the position of strain A3.

**Fig 7.**
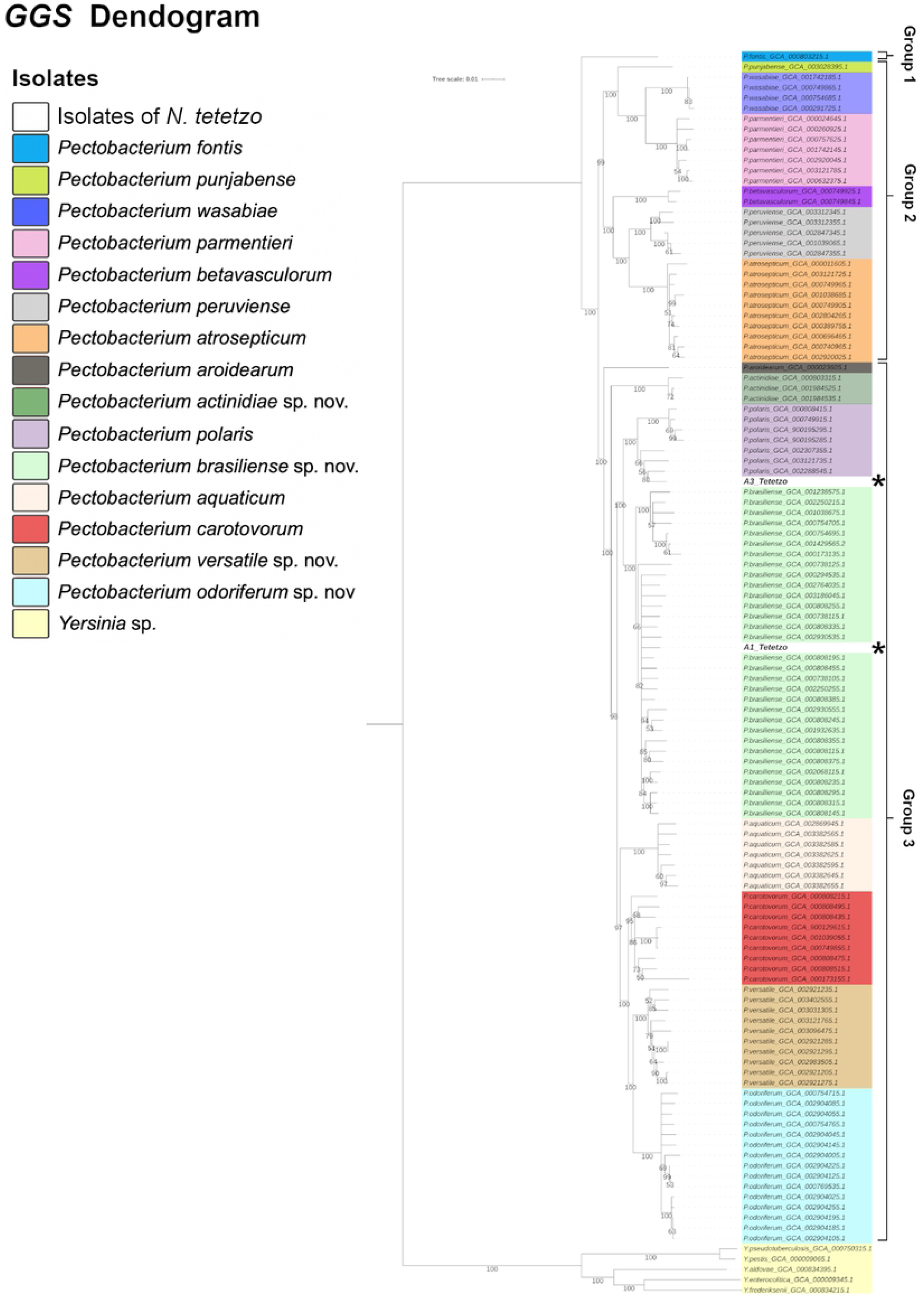
Genomic Similarity Score (GSS) matrix plotted as Neighbor-Joining tree. This tree shows the relationship between isolates A1 and A3 and other *Pectobacterium* species from the comparison of the entire proteome. *Yersinia sp*. Was included as an external group. The consensus tree was inferred from 1,000 replicates. The asterisk (*) indicates the position of strains A1 and A3.

**Fig 8.**
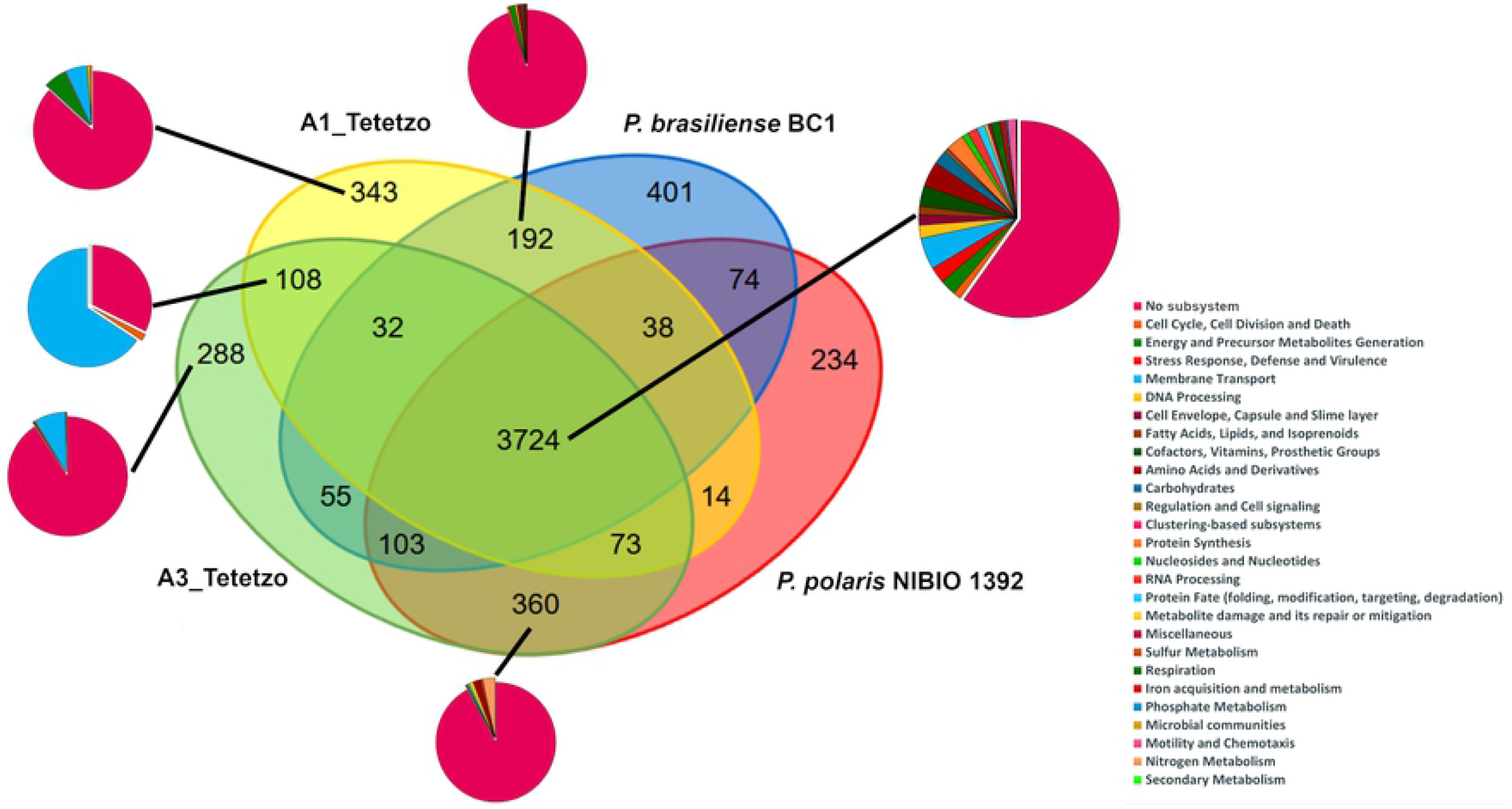
Comparison strains A1 and A3 with two *Pectobacterium* reference genomes. Venn diagram representing the whole genome comparison of sequenced *Pectobacterium* strains A1 and A3, and two complete reference genomes (*P. brasiliense* BC1 and *P. polaris* NIBIO1392). Numbers indicate shared coding sequences (CDS) between strains. Pie charts plot the number of CDS assigned to a main functional role category (subsystem), hypothetical proteins and unknown or unassigned sequences.

### *N. tetetzo* microbiome in health and disease

Finally, we investigated other bacteria that could be participating in the disease course of *N. tetetzo* besides *Pectobacterium*, as it is known for other hosts that as the infection progresses, the microbial community changes. Thus, we analyzed the bacterial community found in an diseased individual and compare it with a healthy cactus. We used metagenomic DNA to describe the microbiome by means of the variable region of the 16S rRNA sequencing. Healthy tissue of *N. tetetzo* contained only three OTUs, assigned at the order level to: Lactobacillales, Oceanospirillales (being the predominant) and Rhizobiales. In comparison, diseased tissue showed greater abundance of OTUs, including: Actinomycetales, Burkholderiales, Caulobacterales, Lactobacillales, Oceanospirillales, Rhizobiales and Sphingomonadales; here, similarly to the healthy sample, the orders Lactobacillales and Oceanospirillales were predominant (Figure 9).

**Figuere 9.**
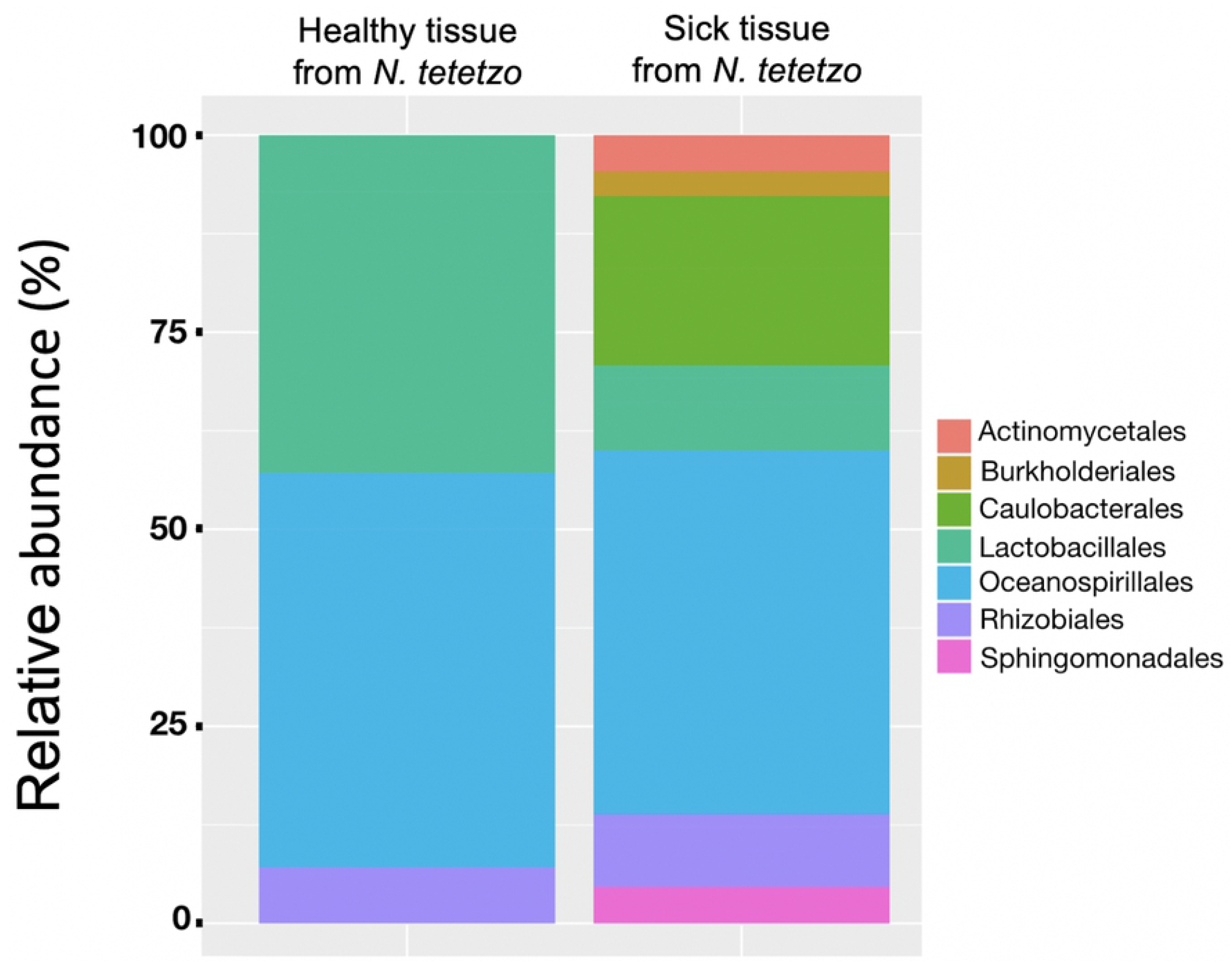
Orders detected in *N. tetetzo* tissue. Three bacterial orders were detected in healthy tissue of *N. tetetzo*: Lactobacillales, Oceanospirillales and Rhizobiales. The orders detected in diseased tissue were: Actinomycetales, Burkholderiales, Caulobacterales, Lactobacillales, Oceanospirillales, Rhizobiales and Sphingomonadales. Healthy tissue is less diverse than diseased tissue.

## Discussion

### Characterization of *Pectobacterium* sp

Here, we give evidence of *Pectobacterium* as the etiological agent of the soft rot of *N. tetetzo*, in agreement with the report of Mejía-Sánchez and collaborators (2019). Biochemical testing indicated identical profiles for three strains isolated from rotten cacti, and the 16S-rRNA gene established that all three belong to the *Pectobacterium* genus. However, further whole genome analyses of A1 and A3, indicated that they are different species: A1 groups with *P. brasiliense* whereas A3 seems to be *P. polaris*, which represents a surprising result. *P. brasiliense* was first reported in Brazil in 2004 by Duarte and collaborators and since then it has reached all continents; meanwhile *P. polaris* was reported in Norway and Poland only recenlty (in 2017). The presence of *Pectobacterium* in different parts of the world, including an isolated area such as Zapotitlán de las Salinas, Puebla is proof that it is a highly mobile ubiquitous pathogen probably present due to globalization. That A1 (and possibly also A8) and A3 are different species could explain the difference on symptom onset that we oberved for each of them. Despite the maceration delay by A3 in comparison to A1 and A8, all isolates caused infection in the hosts analysed: celery and three types of cacti, *O. ficus-indica, N. tetetzo* and *P. pringlei*, adding these to the list of plants susceptible to *Pectobacterium* infection. To our best knowledge, this would represent the first report of *P. polaris* capability for infecting cacti, besides potato [35]. These differences found in maceration capacity are mainly due to the fact that they are two different species according to our phylogenetic analyzes, ANI and GSS, where isolate A1 is grouped with *P. brasiliense* and isolate A3 with *P. polaris*; the ANI pattern agrees with the data presented by Narváez-Barragán and collaborators 2020, while our GSS tree presents a topology similar to that presented by Portier and collaborators (2019). The main advantage of GSS is that it uses both core and pan-genomic information to estimate relatedness between strains. Proteins are the preferred choice to find homologs with large evolutionary distances [37], allowing a better resolution when working with genres.

*Pectobacterium* sp. has also been reported as the main etiological agent of rot in commercial crops in Baja California of prickly pear cactus, *O. ficus-indica* [38]. This is important because Mexico is the main producer and consumer of *O. ficus-indica*, since around 72,000 ha of the country are cultivated to produce fruit and approximately 10,500 ha are cultivated for vegetable production purposes, which is carried out in most of the country’s states [39,40], knowing the identity of the pathogen in these crops is essential for establishing adequate sanitary measures to minimize crop losses.

Few articles have reported the presence of *Pectobacterium* in wild plants, in this case *N. tetetzo, P. pringlei* (although no symptoms have been observed in the field) and in *Carnegiea gigantea* [41], which indicates that we know very little of the ecological function of this pathogen in arid areas where these columnar cacti are found.

### *N. tetetzo* microbiome in health and disease

Finally, our analysis of the bacterial community composition during infection of *N. tetetzo* allowed the identification of different OTUs, however *Pectobacterium* was absent at this disease stage. This was surprising but not unexpected, as infections are dynamic processes in which drastic changes of the bacterial community can occur, specially in long-term infection such that observed in *N. tetetzo*. In support to our observations, Kõiv and collaborators (2015) examined the changes of the endophytic bacterial community in potato tuber (*Solanum tuberosum*) in response to infection by *P. atrosepticum* finding that eight days after inoculation, the Enterobacterales order either decreases or disappears, and others with cellulolytic/pectinolytic capacity such as the Clostridiales appear. This report indicates that although *P. atrosepticum* initiates the infection, it provides a niche for the potato endophytes that then contribute to the degradation of different components of the plant, many of them being opportunists. This seems to be the case also for *N. tetetzo*, which in the absence of infection supports a limited community including the orders Lactobacillales, Oceanospirillales and Rhizobiales. In agreement with our observations, Lactobacillales have been reported in the stem endosphere of other cacti, such as *Myrtillocactus geometrizans* and *Opuntia robusta* [43]. Oceanospirillales are generally halophilic marine bacteria, but their presence could be the result of the complex history of the Tehuacán-Cuicatlán valley, which includes deposition events of salt-rich marine sediments during the lower Cretaceous. However, at more advance disease stages the community diversifies including OTUs from the orders: Actinomycetales, Burkholderiales, Caulobacterales and Sphingomonadales, of which the Actinomycetales stand out as being decomposers of organic matter, contributing to the rot of *N. tetetzo*. Another important order is Burkholderiales since some of these can be pathogens in plants, including species such as *Burkholderia* and *Ralstonia*, in addition to being natural inhabitants of the soil [44]. Caulobacterales have been reported in very diverse habitats, mainly aquatic, in the soil and on the surfaces of eukaryotes, they even form biofilms that allow them to withstand adverse periods, it has also been reported that these bacteria are the main degraders of macroscopic organic aggregates and the release of amino acids to be consumed by other bacteria [45,46], thus probably participating in the rot.

## Conclusions

Isolates A1 and A8 were more virulent with respect to isolate A3, because they are different species. In the same way, the presence of *Pectobacterium brasiliense* and another species of *Pectobacterium, Pectobacterium polaris* in the soft rot of *N. tetetzo* was confirmed.

The genome of isolate A1 (*P. brasiliense*) presented a percentage of guanin-cytosine (%GC) of 52.31, 4,013 coding regions, N50 of 46,958 and an approximate genome size of 4,500,321 bp, while the genome of isolate A3 (*P. polaris*) presented a %GC of 51.96, 4,334 coding regions, N50 of 59,089 and an approximate genome size of 4,873,548 bp.

The microbiome analysis revealed three main OTUs in healthy tissue from *N. tetetzo*: Lactobacillales, Oceanospirillales, and Rhizobiales. Also we detected four OTUs in diseased tissue other than healthy: Actinomycetales, Burkholderiales, Caulobacterales, and Sphingomonadales.

## Acknowledgements

The first author would like to thank Consejo Nacional de Ciencia y TecnologÍa for the Grant (number 816227) that supported his Master degree, and Posgrado de Ciencias Biológicas of Universidad Nacional Autónoma de México (UNAM).

## Funding Information

This project was financed by grants from DGAPA-PAPIIT (IN216317) and CONACYT (A1-S-33379) to R .T.A. And DGAPA-PAPIIT (IN211019) and CONACYT Ciencia Básica (252551) to C. M.-A.

## Supporting information

**S1 Figure. Circular representation of the genomes of Pectobacterium strain A1 and A3**. Concentric circles represent genomic data, from the outermost circle to the innermost: size in Mbp, Contigs, forward strand predicted Coding sequences (CDS), reverse strand CDS, predicted antibiotic resistance genes, virulence genes, transporter genes, G-C content, G-C skew, and unmerged gene clusters identified by PATRIC suite. Contigs were aligned and reordered using the complete genome of type strain *P. carotovorum subsp. brasiliensis* BC1 as a reference for strain A1 and the strain *P. polaris* NIBIO1392 as a reference for strain A3. Those contigs unable to align were placed at the end (red bar).

(TIF)

**S1 Table. Sequences used in this study**

(XLSX)

**S2 Table. ANI identity values**. The green rows indicate *Pectobacterium brasiliense*, the purple rows indicate *Pectobacterium polaris*, the yellow ones indicate *Pectobacterium carotovorum*, the blue column indicates isolate A3_Tetetzo and the red one indicates isolate A1_Tetetzo.

(XLSX)

